# Consideration of genetic and sex effects in mice enhances consilience with human addiction studies

**DOI:** 10.1101/2020.02.14.949784

**Authors:** Michael C. Saul, Jared R. Bagley, Lauren S. Bailey, Udita Datta, Price E. Dickson, Rainy Dodd, Leona H. Gagnon, Spencer B. Hugett, Violet M. Kimble, Michael Leonardo, Sam-Moon Kim, Ashley Olson, Tyler Roy, Sarah A. Schoenrock, Troy Wilcox, J. David Jentsch, Ryan W. Logan, Colleen A. McClung, Rohan H. C. Palmer, Vivek M. Philip, Laura G. Reinholdt, Stacey J. Sukoff Rizzo, Lisa M. Tarantino, Elissa J. Chesler

## Abstract

Concerns about external validity of rodent models and translation of findings across species are often based on narrow investigations of populations with limited diversity. Sources of individual variation – including genetics and sex – are only infrequently encompassed in model organism studies. As with most complex diseases, risk for cocaine use disorder is subject to considerable inter-individual variation. Explicit inclusion of individual differences in rodent research may reveal conserved phenotypes and molecular systems relevant to human addiction. We surveyed cocaine-related traits in both males and females of eight inbred mouse strains whose genomes collectively capture 90% of the genetic diversity of the mouse species. Across these strains, individual differences explained a substantial proportion of variance in cocaine-responsive or cocaine response-predictive behavioral and physiological phenotypes. Wild-derived mouse strains often extended the phenotypic ranges of these behaviors beyond what is observed in conventional laboratory strains. Striatum transcriptional responses to cocaine were also highly dependent upon strain and sex differences; most cocaine-responsive genes were differentially expressed in a manner moderated by strain, sex, or their combination. We compared the strain- and sex-mediated transcriptional responses to cocaine in mice to transcriptomic analysis of people with cocaine use disorder and found that mouse similarity to humans was highly dependent upon mouse genetic background and sex. Specifically, male WSB/EiJ mice and female NOD/ShiLtJ mice exhibited the greatest degree of neural transcriptional consilience with humans with cocaine use disorder. Model organism diversity thus represents a crucial source of biological information that can substantially improve external validity of neuropsychiatric research.

**Significance Statement:** Laboratory mice are widely used in research on neurobiological mechanisms of addiction, but most studies use a single strain and often sex of mice. To assess how individual differences in mice modulate addiction-related traits and how this impacts comparative analysis with humans, we studied cocaine-relevant behaviors and brain molecular correlates in both males and females of genetically diverse mouse strains. In this population, individual differences related to sex and/or genetics explain large proportions of differences in cocaine-related traits. Importantly, brain gene expression data demonstrated that some strains mimic human genomic states more readily than others. Individual differences thus represent a crucial and underdeveloped source of biological information about addiction mechanisms that may influence the translational utility of such studies.

## Introduction

Complex diseases such as neuropsychiatric disorders are typically characterized by the contributions of many genes (Hyman, 2018). Behavioral genetics in non-human animals aims to establish consilience with psychiatrically relevant human systems by studying conserved behaviors and their conserved underlying neural molecular substrates. However, despite mounting evidence of deep conservation in the complex gene systems driving behavior (Saul et al., 2019a; Sinha et al., 2020; Young et al., 2019), uncertainty about the psychiatric relevance of non-human animal systems abounds in the psychiatric genetics community (National Advisory Mental Health Council Workgroup on Genomics, 2018). These doubts likely arise from longstanding and well-characterized difficulties in replicating non-human animal behavioral findings (Crabbe et al., 1999) as well as the failure to translate psychiatric genetics results from any species into the clinic (Hyman, 2012). A critical reassessment of non-human animal behavioral genetics is needed for non-human animals to contribute to the human psychiatric literature.

To simplify experimental design, rodent behavioral and genomic studies of behavioral traits, though often quite detailed in their behavioral and biological scope, most often focus their efforts on identifying individual differences within a single inbred mouse strain for practical reasons (Pascoli et al., 2018; Walker et al., 2018) or within outbred rat populations that often ship from multiple vendors (Fitzpatrick et al., 2013), each with their own genetic bottlenecks. Sex differences are frequently ignored in both human and non-human animal research (Datta et al., 2020). Experimental methodologies agnostic to these known sources of individual differences can produce valuable biological insights in some circumstances, but the omission of sex and genetic variation in these studies limits generalizability without necessarily reducing experimental noise (Prendergast et al., 2014; Tuttle et al., 2018). Further, using a single inbred genetic background can suppress behavioral effects observed in even highly penetrant knockout alleles (Sittig et al., 2016). On the other hand, systematic inclusion of tractable genetic diversity may allow rodents to not just better emulate clinically relevant characteristics, but to contribute new genetic paradigms of understanding neuropsychiatric disorders (Neuner et al., 2019). These important advances in model organism genetics are often ignored in discussions of how to best model neuropsychiatric phenotypes in non-human animals (e.g. Nestler and Hyman, 2010).

As highly prevalent behavioral disorders, substance use disorders drive a public health crisis associated with substantial morbidity and mortality. Illicit substance use disorders afflict approximately 1 in 14 young adults in the United States (SAMHSA, 2017) and drug overdoses are now the leading cause of accidental death among American adults under 55 (Kochanek et al., 2017). Genetic variation and sex differences are both known to influence addiction vulnerability; cocaine use disorder is highly heritable (H^2^ ≈ 0.71) (Goldman et al., 2005) and substance use behaviors show sex differences in both humans and other animals (Becker et al., 2012; Becker and Chartoff, 2019). Consequently, the neurobiology underlying addiction cannot be fully understood without consideration of genetic background and sex in both humans and non-human animals.

Given the genetic diversity in the population of inbred mouse strains (Beck et al., 2000) and the multitude of derived mouse recombinant inbred and heterogeneous stock populations useful for genetic mapping and trait correlation work (Chesler et al., 2005; Logan et al., 2013; Philip et al., 2011), it is possible to sample over diverse genotypes and identify strains and sexes that best mimic the human disease state. Such a strategy can improve translational relevance while surveying these populations for heritable drivers of unidentified genetic mechanisms of disease susceptibility. Though rodent genetic variation does not capture precise human variants, it can be exploited to determine underlying mechanisms in addiction-relevant processes (Bogenpohl et al., 2017; Huggett et al., 2020; Palmer et al., 2019). As a further benefit, genotypic and phenotypic precision, along with high minor allele frequencies and in many cases, a well-randomized population structure, allows genetics studies in rodents to be performed at orders of magnitude lower cost than human GWAS.

To assess the influence of genetic variation, sex, and their potential interactions on cocaine-related phenotypes in mice, we undertook a large-scale evaluation of behavioral, physiological, and brain transcriptomic measures in both male and female mice from the eight inbred founder strains of the Diversity Outbred (DO) mouse heterogenous stock (Saul et al., 2019b). Because these strains include wild-derived inbred strains from the three dominant subspecies of mice, their genomes together capture approximately 90% of the genetic diversity in the species *Mus musculus* (Roberts et al., 2007). We further assessed how strain and sex differences drive consilience with human addiction, identifying which strains and sexes capture significant overlapping brain molecular correlates with human cocaine use disorder.

In the eight founder strains for the DO, we surveyed behavioral and physiological correlates of vulnerability to cocaine use – including multiple novelty response behaviors, circadian molecular rhythm phenotypes, and reversal learning as a measure of reward learning and impulsivity. We directly measured cocaine-related behaviors such as initial locomotor sensitivity to cocaine and intravenous self-administration (IVSA) of cocaine. Finally, using RNA sequencing (RNAseq), we measured the striatum transcriptome response to cocaine in all eight founder strains and compared them to a meta-analysis of postmortem brain tissue from cocaine use disorder patients. Our work represents the first time that many of these behavioral, physiological, and molecular traits have been studied using methods powered to detect sex and genotype effects and the extent to which these interact to moderate one another’s effects.

## Results and Discussion

We first examined heritability, sex differences, and genetic differences moderated by sex among the founders of the DO. Significant effects are reported in the text; all tests are reported in **Supplementary Table S1**.

Response to novelty predicts psychostimulant addiction-related phenotypes in both humans (Ersche et al., 2010) and mice (Dickson et al., 2015). We first assessed differences in behavioral traits related to exploration and response to novelty using the open field, light-dark box, hole board and novel place preference tests. Though novel place preference heritability was moderately weak (H^2^ = 0.16), the other three phenotypes displayed strong to very strong heritability (H^2^ = 0.42-0.55, **Figure 1A-D**). Further, for the phenotype of total entries in the hole board test, an exploratory behavior, there was a significant sex difference (F_1,597_ = 10.69, p = 0.0014), which was apparently driven by lower hole board exploration in males than females for all strains but PWK/PhJ. We detected strain-by-sex interactions in the total entries in the hole board (F_7,597_ = 2.74, p = 0.0083) and in the proportion of distance traveled in the center of the open field (F_7,684_ = 2.09, p = 0.043), indicating that sex differences in these novelty response traits are moderated by genetic background. Total entries in the hole board (**Figure 1A**) and novel place preference (**Figure 1B**) exhibited extended range due to the inclusion of wild-derived strains, demonstrating that wild-derived genetic variability functions in defining an expanded phenotypic range (Wahlsten et al., 2003). Transitions between the sides of the light-dark box also showed phenotypes toward the extreme in wild-derived mouse strains along with NOD/ShiLtJ mice (**Figure 1D**).

**Figure 1:**
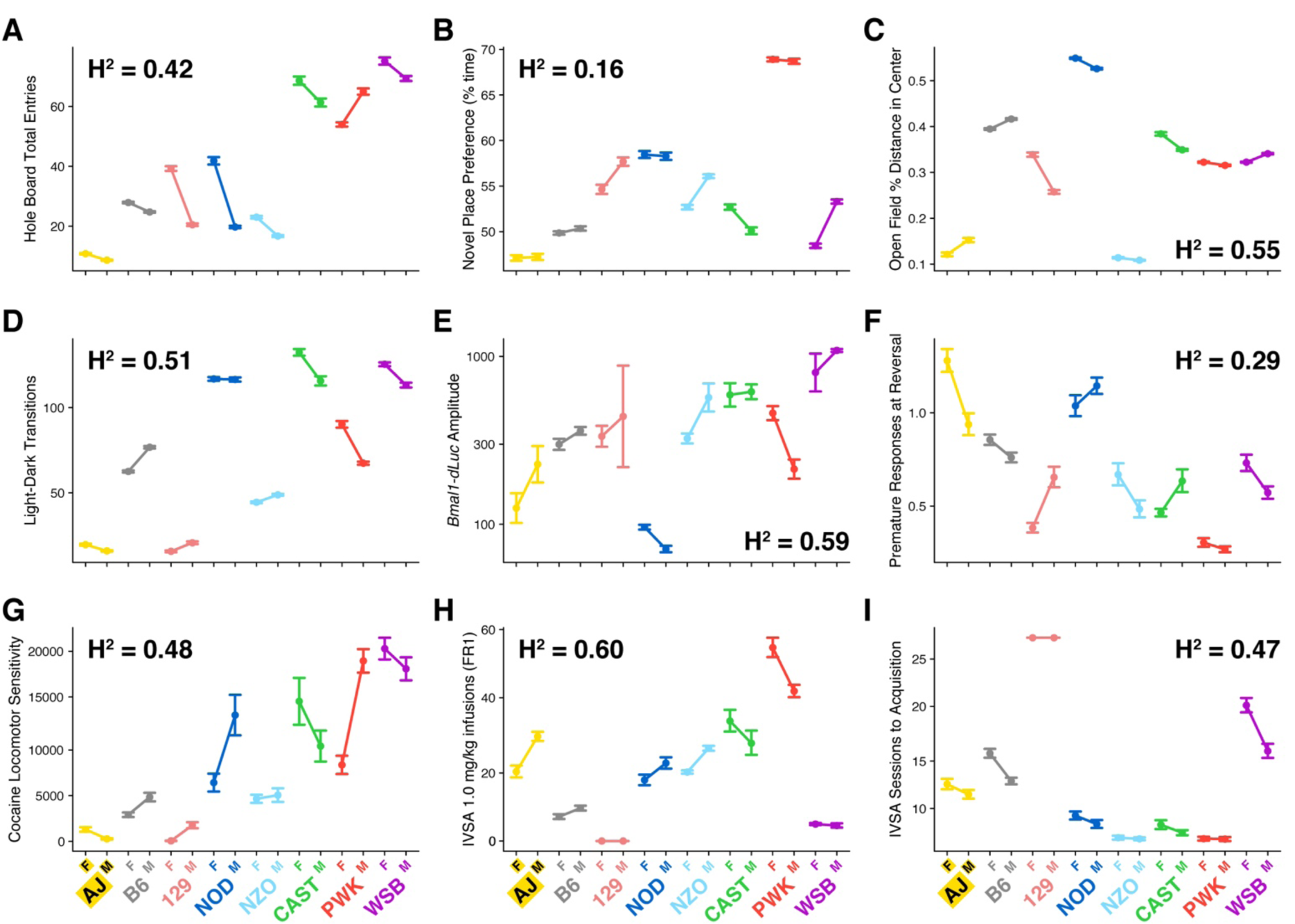
Heritable differences in cocaine-related behavioral and physiological traits. All plots show the mean ± the standard error in both sexes with females on the left. **A)** Hole board total entries, **B)** Novel place preference (% time in novel zone), **C)** Open field (% time in center of open field), **D)** Transitions between light and dark in the light-dark box, **E)** Circadian rhythm fibroblast *Bmal1-dLuc* luminescence amplitude, **F)** Reversal learning premature responses to the hole rewarded during acquisition, **G)** Initial locomotor sensitivity to cocaine (cm moved on Day 3 – Day 2), **H)** Cocaine intravenous self-administration number of infusions at 1.0 mg/kg FR1, and **I)** Sessions to acquisition of cocaine intravenous self-administration.

Circadian rhythm and reward-related behavioral phenotypes are co-inherited and rhythm disruptions are linked to development and progression of substance use disorders (Logan et al., 2014). Further, the molecular clock system directly influences the expression of dopamine receptors in the striatum involved in the modulation of cocaine reward-related behaviors (Ozburn et al., 2015). A cell-based assay on fibroblasts derived from each of the founder strains in which a *Bmal1-dLuc* reporter was utilized for circadian measurement of luciferase bioluminescence to assess differences in circadian rhythmicity (Kim et al., 2016; Ramanathan et al., 2014). We found very high heritability of the amplitude of these rhythmic patterns (H^2^ = 0.59, **Figure 1E**), but no significant sex differences or strain x sex interactions. These results suggest that genetic differences in the molecular clock are one potential mechanism for individual differences in addiction-related phenotypes.

Reversal learning tasks evaluate impulsive and compulsive behaviors that predict addiction liability (Izquierdo and Jentsch, 2012). Within reversal learning paradigms, one measure of impulsivity is the number of premature responses produced during the reversal phase of the task. This phenotype is similar to an impulsivity-related measure implemented in the five choice serial reaction time task (Dalley et al., 2007). Premature reversal responses in our study showed moderate heritability across our eight strains of mice (H^2^ = 0.29, **Figure 1F**), but no significant sex differences or strain x sex interactions. We observed the lowest rate of premature responding in the wild-derived PWK/PhJ strain.

We next sought to assess individual differences in initial locomotor response to cocaine, a behavioral phenotype that predicts subsequent drug use in humans (de Wit and Phillips, 2012). In these diverse mouse strains, initial cocaine sensitivity showed strong heritability (H^2^ = 0.48 for initial sensitivity, **Figure 1G**). Wild-derived strains, particularly WSB/EiJ and PWK/PhJ, exhibited the highest initial sensitivity to cocaine.

Operant drug self-administration procedures directly quantify reinforced responding for drug (Dickson et al., 2015) and produce profound transcriptional responses in inbred strain (Walker et al., 2018). Heritability of cocaine IVSA traits in the founders of the DO was strong (H^2^ = 0.47 for sessions to acquisition of IVSA, **Figure 1H**) to very strong (H^2^ = 0.60 for total infusions at FR-1 1.0 mg/kg, **Figure 1I**). The high heritability of infusions earned is similar in magnitude to the observed heritability of human cocaine use disorder (Goldman et al., 2005). Of note, 129S1/SvImJ mice do not acquire IVSA and do not take any cocaine during the acquisition phase. The largest phenotypic range manifests in wild-derived mice; PWK/PhJ mice acquired IVSA behavior very quickly and self-administered the most infusions of cocaine at FR-1 1.0 mg/kg – about four times as much as C57BL/6J – while WSB/EiJ mice self-administered the fewest infusions of cocaine relative to the other strains that do acquire IVSA behavior. The broad phenotypic range of volitional cocaine-taking behaviors in these genetically diverse mouse strains represents a clear opportunity to study mechanisms underlying variation in the initiation of cocaine of addiction.

Because addiction-related phenotypes are highly heritable and sometimes exhibit sex differences moderated by genetics, we next sought to assess heritable differences in the molecular response to cocaine. Male and female mice of the eight DO founder strains were given multiple injections of cocaine or saline over 19 days. Samples for bulk RNAseq were collected from striatum 24-48 hours after the final injection of repeated administration of either cocaine or vehicle (sham). First, we documented that in the absence of cocaine, most transcripts exhibited expression that was moderate to strong at baseline in the founder strains (median H^2^ = 0.29) and that subspecies of origin explains the greatest amount of variance in expression (**Figure 2A**). Using a linear modeling approach (Chen et al., 2014; Phipson et al., 2016), we found few expression differences attributable solely to the effect of cocaine treatment. Instead, significant effects of cocaine arose in interaction with individual differences such as genetic background, sex, and their interaction. The greatest quantity of differentially expressed transcripts was observed in the strain-by-sex-by-drug treatment three-way interaction (1,282 genes at q < 0.01, **Figure 2B, Supplementary Table S2**). Some strains showed stronger cocaine effects as measured by number of differentially expressed genes than others; 129S1/SvImJ and PWK/PhJ had many genes whose expression is influenced by cocaine and sex-by-cocaine interactions (PWK/PhJ: 89 genes at q < 0.01, **Figure 2C**) while the commonly used C57BL/6J strain had very few genes influenced by cocaine (four genes at q < 0.01, **Figure 2C**). Because statistical power was approximately equal for all strains involved in this study, these differences likely reflect real individual differences in the brain’s sex-specific responses to cocaine.

**Figure 2:**
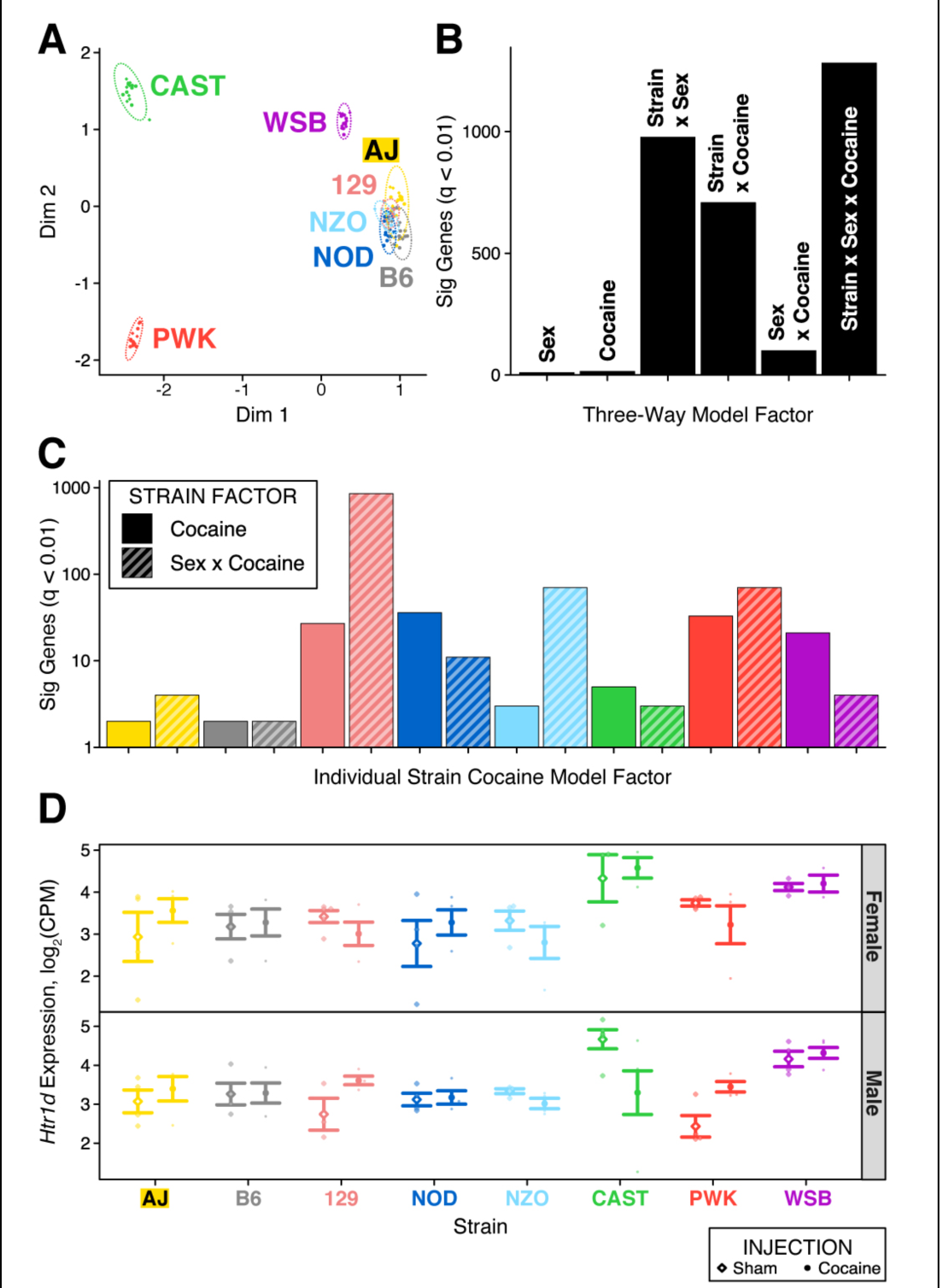
Individual differences in striatum cocaine response transcriptomes. **A)** High heritability of baseline transcriptome partitions the subspecies in a multidimensional scaling plot. Colors in Panel A carry forward to other figure panels. **B)** In a three-way analysis of strain, sex, and cocaine, many genes showed significant expression changes in response to cocaine in interaction factors; the strain x sex x cocaine interaction factor identified 1,282 DEGs. In contrast, there were very few significant DEGs in the main effect of cocaine. **C)** Two-way models show that the magnitude of transcriptional responses to cocaine differs by strain. B6 has few differentially expressed genes while 129 and PWK have many. The y-axis is log_10_. **D)** The serotonin 1D receptor gene *Htr1d* is differentially expressed in males of 129S1/SvImJ, PWK/PhJ, and CAST/EiJ strains. CAST/EiJ males show the opposite directionality of differential expression to PWK/PhJ and 129S1/SvImJ males.

This genetics- and sex-inclusive approach identified many more changes in gene expression following repeated cocaine exposure than had we used only a single sex in a single inbred strain. For instance, we identified differential expression of the serotonin receptor 1D gene *Htr1d* in the strain-by-sex-by-cocaine interaction factor. This gene, while not altered in C57BL/6J animals, is upregulated in males of 129S1/SvImJ and PWK/PhJ strains, but downregulated in males of the CAST/EiJ strain (**Figure 2D**).

Some of the heritable transcriptome differences we observed corroborate previous work showing heritable differences in pharmacokinetics of cocaine (Wiltshire et al., 2015) – for example, PWD/PhJ mice – closely related to the high cocaine taking PWK/PhJ strain – are known to have some of the highest brain concentrations of cocaine shortly after injection. These pharmacokinetic differences do not account for the strong differences between strains such as 129S1/SvImJ and A/J, whose brain cocaine pharmacokinetic profiles are very similar (Wiltshire et al., 2015). Further research on highly diverse mice may resolve the mechanisms driving individual differences between strains with closely matched pharmacokinetic profiles.

We assessed to what degree strain and sex combinations mimic human genomic findings to understand how individual differences’ influence on consilience with human research. We used the MSET method (Eisinger et al., 2013) to identify statistically significant overlaps of orthologous genes between a human cocaine use disorder brain transcriptome dataset and analogously analyzed cocaine response transcriptomes within each individual strain and sex combination (Huggett et al., 2020). The magnitude of overlap was highly variable; the strongest overlaps were in NOD/ShiLtJ female and WSB/EiJ male mice, though the NOD/ShiLtJ female finding is not robust to different significance thresholds and does not survive stringent multiple testing correction (**Figure 3A, Supplementary Table S3**. Other strains, including the most widely used C57BL/6J, do not strongly mimic cocaine use disorder brain transcriptomes, a finding consistent with previous observations of C57BL/6J males (Huggett et al., 2020). When we examined the directionality of differential expression overlaps in NOD/ShiLtJ females and in WSB/EiJ males, we found concordance in the upregulated and downregulated genes between human and mouse data; in this experiment, WSB/EiJ males mostly captured downregulated aspects of expression while NOD/ShiLtJ females mostly captured upregulated aspects (**Figure 3B**). Genes within the NOD/ShiLtJ female set include directional matches in immediate early genes such as *Arc*, and *Junb* while genes downregulated in WSB/EiJ males include multiple oxidative phosphorylation-related genes such as *Ndufa5, Cox6c2*, and *Ndufaf2*. The presence of two mitochondrial Complex I subunit genes is particularly interesting; cocaine treatment disrupts Complex I in the rat brain (Cunha-Oliveira et al., 2013) and a Complex I subunit gene is associated with cocaine use disorder in humans (Huggett and Stallings, 2020a).

**Figure 3:**
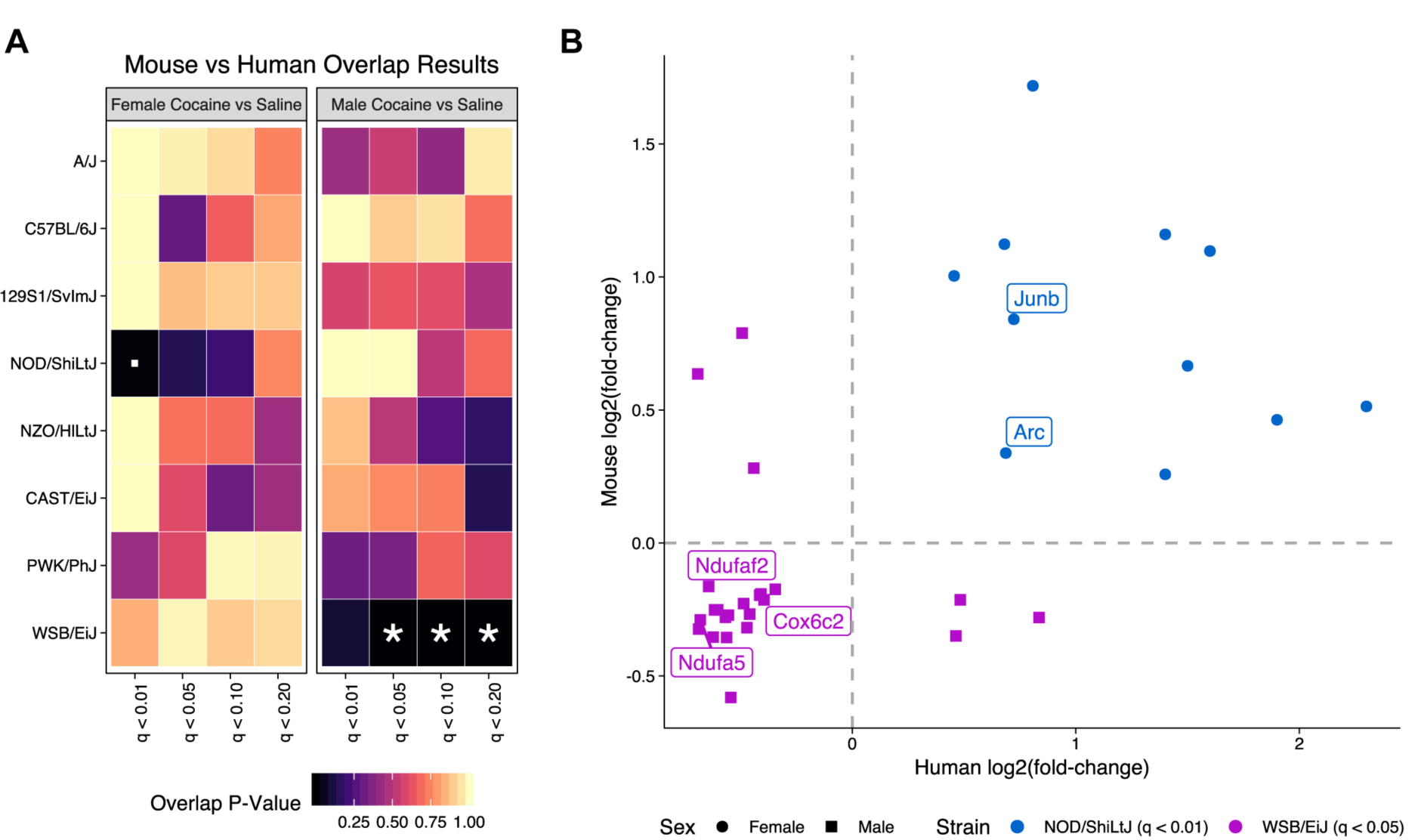
Consilience differs by strain and sex. **A)** According to MSET p-values, a measure of higher-than-expected overlap, NOD/ShiLtJ female (left) and WSB/EiJ male (right) transcriptome differential expression shows statistically significant overlap with the union of multiple human cocaine use disorder brain transcriptome differential expression sets (multiple testing correction significance: ▪ = q < 0.20, * = q < 0.05). **B)** Overlapping gene lists show concurrent directionality. NOD/ShiLtJ female mice display increased expression of immediate early genes while WSB/EiJ males display decreased expression of oxidative phosphorylation genes.

To describe the molecular systems dysregulated after repeated cocaine exposure in these strains, we performed separate Gene Ontology Biological Process (GO BP) analysis of the genes shared between human cocaine use disorder and WSB/EiJ male or the NOD/ShiLtJ female differentially expressed gene sets. WSB/EiJ male differentially expressed genes were enriched mostly for RNA binding and mRNA processing while NOD/ShiLtJ female differentially expressed genes were enriched for nucleobase-related metabolic processes (**Supplementary Table S5**. Together, these results imply that similarities mostly capture alterations in transcriptional regulatory components.

Because the transcriptome results derive from the same group of mice as cocaine locomotor sensitivity, it is of particular note that WSB/EiJ mice show the strongest behavioral response in cocaine-induced locomotor sensitivity of all the strains tested (**Figure 1G**). Variants that alter initial sensitivity to drugs are among the best supported and described addiction candidates in humans, for example, nicotinic acetylcholine receptor polymorphisms associated with smoking alter the receptor’s sensitivity to nicotinic agonists (Bierut et al., 2008). This result implies that WSB/EiJ male mice experience human-relevant alterations in their neurobiological states due to differential cocaine sensitivity after repeated cocaine injection. However, WSB/EiJ mice take fewer infusions of cocaine at FR-1 1.0 mg/kg than any other strain that reliably acquires IVSA (**Figure 1H**). Though the finding of higher consilience associated with lower cocaine-taking behavior appears to contradict the conventional interpretation of cocaine self-administration, where higher cocaine intake indicates higher addiction vulnerability (Piazza et al., 2000), we note that experimenter-administered cocaine affects neural signaling differently than self-administered cocaine (McCutcheon et al., 2011). Consequently, this finding should demonstrate that individual differences are highly important to consider for translational research using non-human animals, but should not be taken as a specific recommendation of WSB/EiJ males as a model for human cocaine use disorder phenotypes.

The translational relevance of non-human animals – often represented by mice – is the subject of substantial discussion (Seok et al., 2013; Takao and Miyakawa, 2015). Within addiction biology, a particularly active source of ferment is the question of how rodents may best recapitulate aspects of addictive behaviors such as compulsive use and reinstatement (Ahmed, 2012). Here, we further this discussion by demonstrating that similarity of rodent molecular systems to those involved in human neuropsychiatric traits are heterogeneous and contingent upon the specific rodents chosen. Within conventional inbred mouse strains, NOD/ShiLtJ females and WSB/EiJ males showed the highest consilience with human cocaine phenotypes in the brain as measured by similarity in genomic response. This finding has broad implications for translational work. In many cases, the study of a single strain and/or only one sex of mice within an experiment may limit the generalizability of the data and diminish potential relevance for human addiction. Rather, it may be advantageous to explicitly take individual differences of the type reported here into account. Strategies for querying diversity include identification of specific strains that exhibit characteristics of the disease-susceptible population at a behavioral or molecular level or surveys of a diverse population to ensure that natural variation is represented, which allows queries of known biological sources of variation in addiction vulnerability.

An important aspect of experimental rigor is the external validity of model organism research paradigms. Here we show that this validity is in part driven by the organism under investigation in addition to other characteristics of the research paradigm. Genetic variation is a valuable resource for the discovery of biological mechanisms of addiction (Kumar et al., 2013; Ruan et al., 2020). Similar to humans, individual differences among mice greatly influence behavioral, physiological, and transcriptomic cocaine-related traits. For many of these traits, individual differences explain a substantial proportion of the variation. This variation can be exploited to enhance consilience of biological systems under study, or to discover the underlying biological mechanisms of vulnerability to disease. Individual variation in addiction-related traits is a largely untapped resource that can be exploited to improve and accelerate discovery of neurobiological and genetic mechanisms related to risk for addiction as well as other complex diseases.

## Methods

### Standard Operating Procedures

All methodologies used in this work are documented in depth in the Center for Systems Neurogenetics of Addiction’s Standard Operating Procedures (SOPs) (https://www.jax.org/research-and-faculty/research-centers/systems-neurogenetics/data-resources). The brief methods appearing below summarize these SOPs and reference specific SOPs.

### Animals

Mice from the following strains were surveyed in these experiments: A/J (JAX stock #000646), C57BL/6J (JAX stock #000664), 129S1/SvImJ (JAX stock #002448), NOD/ShiLtJ (JAX stock #001976), NZO/HlLtJ (JAX stock #002105), CAST/EiJ (JAX stock #000928), PWK/PhJ (JAX stock #003715), and WSB/EiJ (JAX stock #001145). These strains are the founders of the DO heterogeneous stock and CC recombinant inbred strains. Surveys of these eight strains can demonstrate statistical heritability patterns that justify further dissection using the derived resources (Saul et al., 2019b). The mice used in the Research Animal Facility at The Jackson Laboratory came from breeding colonies maintained in the Research Animal Facility. These colonies were derived from production colonies at The Jackson Laboratory and breeders were replaced with animals from The Jackson Laboratory’s production colony at least every five generations. The mice used in the Jentsch Lab at Binghamton University were shipped to the Jentsch Lab from either the Research Animal Facility colonies or from production colonies in The Jackson Laboratory. The studies described utilized a total of 1,085 mice (**Supplementary Table S4**). All procedures were approved by the Jackson Laboratory of Mammalian Genetics or Binghamton University institutional animal care and use committees.

### Mouse Husbandry and Housing

Mice were housed according to the CSNA animal housing SOP (https://www.jax.org/-/media/jaxweb/files/research-and-faculty/tools-and-resources/system-neurogenetics/csna-animal-housing.pdf).

### Cocaine

Cocaine hydrochloride was provided by the National Institute on Drug Abuse Drug Supply Program Division of Therapeutics and Medical Consequences (catalog number: 9041-001). Cocaine was stored in powder form at room temperature until it was formulated into 0.9% Saline (100 mg/mL clear solution) in various concentrations specific to each experiment according to their individual SOPs.

### Novelty Response Behavioral Phenotypes

Open field, light-dark box, hole board, and novel place preference behavioral paradigms were conducted in this order on consecutive days over the course of a week during the light phase of the light:dark cycle. Open field data were collected for 60 minutes according to the SOP (https://www.jax.org/-/media/jaxweb/files/research-and-faculty/tools-and-resources/system-neurogenetics/open-field-assay.pdf?la=en&hash=32DDAFF2B17B2D4961C136C5616C4982AC23EC3B). Light-dark data were collected for 20 minutes with the mouse starting in the light side of the chamber facing the dark side according to the SOP (https://www.jax.org/-/media/jaxweb/files/research-and-faculty/tools-and-resources/system-neurogenetics/light-dark-assay.pdf?la=en&hash=A63CF8D22EB7936CF6C69A3178373981F4016675). Hole board data were collected for 20 minutes according to the SOP (https://www.jax.org/-/media/jaxweb/files/research-and-faculty/tools-and-resources/system-neurogenetics/hole-board-assay.pdf?la=en&hash=EC343A797D37209CF64D34E6031608A511D8E15D). Novel place preference included a five minute acclimation period to a center chamber, a 10 minute exposure period to a randomized exposure side, a five minute acclimation period, and a final test period consisting of a five minute habituation period again to the center and a 20 minutes preference assessment for which both the novel side and the initial familiar exposed side were accessible according to the SOP (https://www.jax.org/-/media/jaxweb/files/research-and-faculty/tools-and-resources/system-neurogenetics/novelty-place-preference-assay.pdf?la=en&hash=B5D2D0FC9028B408E84729C0C8832C580AB8E039). All mice from the Center for Systems Neurogenetics of Addiction were tested through this novelty pipeline prior to any other test and were then randomized and assigned into either reversal learning, cocaine locomotor sensitization, or cocaine intravenous self-administration. The novelty study produced observations from a total of 783 mice.

### *Bmal1-dLuc* Circadian Rhythm Data

Data for circadian rhythm were measured in primary fibroblast cultures generated from skin biopsies in the founders. To isolate fibroblasts, ear biopsies (one mm in diameter) were digested in Dulbecco’s Modified Eagle’s Medium (DMEM, HyClone) containing 2.5 mg/ml collagenase D (Gibco) and 1.25 mg/ml pronase (Millipore) for 90 mins and then plated in DMEM growth media containing 10% Fetal Bovine Serum (FBS, HyClone), 292 μg/ml L-glutamine (HyClone), 100 units/ml penicillin (Hyclone) and 100 μg/ml streptomycine (HyClone). *Bmal1-dLuc* reporter was delivered to fibroblasts by lentiviral-mediated gene delivery (VectorBuilder). Following synchronization of rhythms by 15 μM forskolin (Sigma) for two hours, the temporal patterns of *Bmal1-dLuc* bioluminescence was recorded for ∼70 seconds at intervals of 10 minutes over six to seven days from fibroblast cultures in DMEM recording media containing 15 μM forskolin, 25 mM HEPES (Gibco), 292 μg/ml L-glutamine, 100 units/ml penicillin, 100 μg/ml streptomycine, and 10 μM luciferin (Promega) by an automated 32-channel luminometer (Lumicycle, ActiMetrics) in a standard tissue culture incubator at 32°C. The amplitude of bioluminescence rhythms was determined from baseline-subtracted data using the damped sine fit and Levenberg-Marquardt algorithm (Izumo et al., 2003). The circadian study produced observations from a total of 56 mice.

### Reversal Learning

Data for reversal learning were collected at both JAX and Binghamton University using the SOP (https://www.jax.org/-/media/jaxweb/files/research-and-faculty/tools-and-resources/system-neurogenetics/reversal-learning-assay.pdf?la=en&hash=8484E47B170462960E11C1FAEEE6FF3CE6FDFC08). The reversal learning data produced observations from a total of 202 mice.

### Initial Locomotor Sensitivity to Cocaine

Data for initial locomotor sensitivity were collected as described previously (Schoenrock et al., 2020) using data from days 1-3 in the SOP for locomotor behavioral sensitization to cocaine (https://www.jax.org/-/media/jaxweb/files/research-and-faculty/tools-and-resources/system-neurogenetics/cocaine-locomotor-sensitization-assay.pdf?la=en&hash=9E5D4C248C3BCCAD947C164AE81663C13A77EB0D). Briefly, mice were placed into the open field arena for 30 minutes, removed, and injected i.p. with either saline (days 1-2) or 10 mg/kg cocaine (day 3) and returned to the open field arena for 60 minutes. Distance moved after injection on day 3 minus day 2 was uses as a measure of initial locomotor sensitivity to cocaine. The sensitization study produced observations from a total of 230 mice.

### Cocaine Intravenous Self-Administration

Prior to cocaine intravenous self-administration, mice were implanted with a jugular catheter and allowed a minimum of 10 days for post-operative recovery. In an operant conditioning paradigm, mice were allowed to acquire cocaine self-administration at 1.0 mg/kg, then evaluated for dose-response effects at eight different doses. After a stabilizing dose at 1.8 mg/kg, extinction-related responses during seven days of withdrawal were recorded. Finally, cued reinstatement was recorded for two days. Self-administration in these eight mouse strains was performed according to v1.0 of the CSNA’s SOP (https://www.jax.org/-/media/jaxweb/files/research-and-faculty/tools-and-resources/system-neurogenetics/intravenous-self-administration-ivsa-paradigm.pdf?la=en&hash=FA64135F219C7DF65937A1CF9270301B0E771836). The intravenous self-administration study produced observations from a total of 217 mice.

### Data Deposit

Data for each phenotype will be deposited in the Mouse Phenome Database (MPD) (Bogue et al., 2019) upon publication.

### Heritability Calculations

For each trait, heritability was calculated from linear models using the isogenic strain as the independent categorical variable using the following equation:

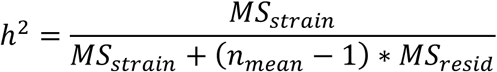

where *MS*_*strain*_ is the mean square of the strain effect, *n*_*mean*_ is the mean number of samples within each strain, and *MS*_*resid*_ is the mean square of the residuals. For the reversal learning data, an additional additive covariate of site was included in the model to account for inter-lab variation. This term was not utilized in the heritability calculation.

For some traits such as number of infusions at FR-1 1.0 mg/kg cocaine self-administration, a single strain such as 129S1/SvImJ showed little to no variation, which may upwardly bias heritability calculations. For these traits, heritability was calculated both with and without the low variance strain. The data reported in the paper rely upon the inclusive calculation, but results of both methods of calculation are reported for completeness (see **Supplementary Table S1**).

### RNAseq

Striatum tissue was collected during the light stage of the light:dark cycle between 24 and 48 hours after the final injection in the cocaine behavioral sensitization protocol according to the SOP (https://www.jax.org/-/media/jaxweb/files/research-and-faculty/tools-and-resources/system-neurogenetics/post-sensitization-tissue-collection.pdf?la=en&hash=9E6CD8DEB39606B791A5D25F6CD0611EF14D96A7). Tissue was collected for both sexes of each founder strain exposed to either sham (saline) or 10 mg/kg IP cocaine.

RNA was isolated from striatum tissue using the MagMAX mirVana Total RNA Isolation Kit (ThermoFisher) and the KingFisher Flex purification system (ThermoFisher). Tissues were lysed and homogenized in TRIzol Reagent (ThermoFisher). After the addition of chloroform, the RNA-containing aqueous layer was removed for RNA isolation according to the manufacturer’s protocol, beginning with the RNA bead binding step.

RNA concentration and quality were assessed using the Nanodrop 2000 spectrophotometer (Thermo Scientific) and the RNA Total RNA Nano assay (Agilent Technologies). 2µl of diluted 1:1000 diluted ERCC Spike-in Control Mix 1 (Ambion by Life Technologies) was added to 100ng of each RNA sample prior to library construction. Libraries were prepared by the Genome Technologies core service at The Jackson Laboratory using the KAPA RNA Hyper Prep Kit with RiboErase (HMR) (KAPA Biosystems), according to the manufacturer’s instructions. Briefly, the protocol entails depletion of ribosomal RNA (rRNA), RNA fragmentation, first and second strand cDNA synthesis, ligation of Illumina-specific adapters containing a unique barcode sequence for each library, magnetic bead size selection, and PCR amplification. Libraries were checked for quality and concentration using the D5000 ScreenTape assay (Agilent Technologies) and quantitative PCR (KAPA Biosystems), according to the manufacturers’ instructions.

RNAseq libraries were pooled and sequenced by Novogene in 150 bp paired-end format on an Illumina NovaSeq 6000 sequencer targeting 90 million read pairs per sample. Sequencing achieved a median read depth of 132 million reads. The resultant reads were determined to be of consistently high quality using fastqc v0.11.3 and MultiQC v1.2.

Reads were generated from raw data and demultiplexed using BCL2Fastq v2.18.0.12, concatenated by sample, and aligned with the STAR aligner v2.6.1 (Dobin et al., 2013) to the GRCm38 mouse reference genome with v94 of the Ensembl transcriptome. Transcript-level quantification was estimated using RSEM v1.3.0 (Li and Dewey, 2011) on a transcriptome BAM file produced as an output of this alignment. The data were imported into R v3.5.1 and summarized to the gene level using tximport v1.10.1 (Soneson et al., 2016), TMM-normalized using edgeR v3.24.3 (Chen et al., 2014), and imported into limma v3.38.3 (Ritchie et al., 2015) using the log_2_-transformation function voom. We compared multivariate approaches modeling with interaction factors between edgeR and voom+limma approaches and found that voom+limma performs better than edgeR for controlling false negatives. Upon initial examination of the findings, we identified intermittent contamination with choroid plexus, which potentially derives from the ventricular aspect of the dorsal striatum. Correcting for this contamination necessited an additive covariate for choroid plexus consisting of log-mean CPM values of *Kl* and *Ttr* expression, unambiguous markers for choroid plexus (Sathyanesan et al., 2012). These values were log_2_ transformed for work in limma. For An overall model for all strains included this choroid plexus factor as a nuisance variable plus the main effects of strain, sex, and cocaine injection and all of their interactions. Individual models included the choroid plexus nuisance variable plus sex, cocaine injection, and sex:cocaine injection interaction. Correction for local false discovery rates utilized the qvalue package in R v2.14.1 (Storey and Tibshirani, 2003). Because brain transcriptional changes are subtle (Hitzemann et al., 2014), all results reported are at q < 0.01 with no fold-change cutoff (**Supplementary Table S2**).

Raw data and transcript-level expression estimates will be deposited in the Gene Expression Omnibus (Barrett et al., 2012) upon publication (accession number: GSEXXXXX).

### Cross-Species Gene List Comparison

To assess the molecular correspondence of mouse cocaine self-administration with human cocaine use, we compared the results of the current study to differentially expressed genes (BH-FDR < 0.05) associated with cocaine use disorder (CUD) in the midbrain (n = 20, 50% CUD, *M*AGE = 49.2, s.d. = 3.9; (Bannon et al., 2014), microarray), hippocampus (n = 15, 46.7% CUD, *M*AGE = 39.4, s.d._AGE_ = 39.4; <u>Huggett and Stallings, 2020a, 2020b</u>, RNA-sequencing) and dlPFC neurons (n = 36, 52.7% CUD, *M*AGE =35.0, s.d._AGE_ = 11.0; (Huggett and Stallings, 2020a); neuron-specific RNA-sequencing). The aforementioned studies utilized methods that maximized power for identifying differentially expressed genes and used case/control analyses that compared individuals with CUD to matched cocaine free controls. A list of all the differentially expressed genes can be found on GeneWeaver (https://www.geneweaver.org/; GS398242).

A key of 1:1 orthologs between humans and mice was generated from the MGI Vertebrate Homology indices (accessed 2020-05-18). This key was used to compare the human gene set to contrast tables within strain and sex combinations. To perform these comparisons, we used the MSET algorithm (Eisinger et al., 2013) as contained in v1.16.6 of the msaul/msaul R package (https://github.com/msaul/msaul). MSET p-values were corrected for multiple comparisons using the qvalue package in R v2.14.1 (Storey and Tibshirani, 2003). Gene Ontology Biological Process (GO BP) analysis on overlapping genes was performed on mouse Ensembl identifiers using the AmiGO v2 web tool (accessed 2020-05-28) on GO Ontology Database doi:10.5281/zenodo.3727280 with Fisher’s exact tests and Bonferroni correction settings (Carbon et al., 2009).

## Supplemental Material

**Supplementary Table S1**: Statistical test results for heritability and genotype-by-sex interactions.

**Supplementary Table S2**: Differential expression results for all tests discussed.

**Supplementary Table S3:** Quantification of overlap significance between human cocaine use disorder transcriptome and individual strain and sex cocaine transcriptomes.

**Supplementary Table S4**: Sample sizes for all strain and sex combinations for studies reported.

**Supplementary Table S5:** Gene Ontology Biological Process results for WSB/EiJ male and NOD/ShiLtJ female overlapping gene sets.

Raw behavioral data will be deposited in the Mouse Phenome Database upon publication. Gene expression data will be deposited in the Gene Expression Omnibus upon publication.

All scripts, code, and metadata used for analysis are deposited in GitHub (repository: github.com/msaul/csna_founders_survey_2020)

## Author Contributions

EJC, PED, LMT, SAS, JDJ, RWL, CAM, LGR, VMP, and SJSR conceived the studies. SJSR, PED, JRB, LSB, SAS, RD, ML, AO, TR, TW, and LHG designed and implemented the behavioral experiments. SMK designed and implemented the circadian transcriptional experiment. JRB, LSB, UD, PED, ML, SMK, AO, TR, SAS, TW, LHG, VMP, and MCS analyzed the behavioral data. SMK, MCS, and VMP analyzed the circadian transcriptional data. MCS, VMK, and VMP analyzed the RNAseq data. MCS, UD, PED, SBH, JRB, LSB, SAS, TW, LHG, VMP, and EJC interpreted the behavioral results. SMK, MCS, and VMP interpreted the results of the circadian transcriptional experiment. MCS, VMK, VMP, and EJC interpreted the RNAseq results. MCS, SBH, RHCP, and EJC conceived, designed, analyzed, and interpreted the comparison between mouse and human gene sets. MCS and EJC wrote the manuscript.

## Acknowledgements

We gratefully acknowledge Genome Technologies service at The Jackson Laboratory for their expert assistance on the RNAseq experiment, Surgical Services at The Jackson Laboratory for their work on the intravenous self-administration paradigm, and the NIDA Drug Supply for providing cocaine. We would further like to thank Stephen Krasinski for critical reading and comments during manuscript preparation, Robert W. Williams for helpful discussion on the analysis, and C. Herbert Pratt for scientific program management. This work was funded by NIH P50 DA039841 (Center for Systems Neurogenetics of Addiction) to EJC, LMT, JDJ, RWL CAM, VMP, LGR, and SJSR as well as NIH R01 DA037927 to EJC, DP1 DA042103 to RHCP, and K99 DA043573 to PED.

